# Predator-mediated interactions through changes in predator home range size can lead to local prey exclusion

**DOI:** 10.1101/2022.09.30.510100

**Authors:** Andréanne Beardsell, Dominique Berteaux, Frédéric Dulude-De-Broin, Gilles Gauthier, Jeanne Clermont, Dominique Gravel, Joël Bêty

## Abstract

The effects of indirect biotic interactions on species occurrence are difficult to quantify in the wild. In theory, the exclusion of a prey species can occur through the numerical and functional responses of a predator to another prey. Few studies assessed the relative effects of these responses on the net interaction strength between multiple prey sharing common predators, in part because empirically based multi-species functional response models are very rare. To investigate whether the presence of a prey species affects predation rates and population growth rate of another prey species, we used a multi-prey mechanistic model of predation along with a population matrix model. The predation model was parameterized using a combination of behavioral, demographic, and experimental data acquired in an arctic vertebrate community. It includes the arctic fox (*Vulpes lagopus*), a predator feeding primarily on small mammals as well as eggs of various bird species such as sandpipers and colonial nesting geese. Our results showed that the positive effects of the presence of a goose colony on sandpiper nesting success (due to the handling time of goose eggs by the predator) were outweighed by the negative effect of an increase in fox density. The numerical response of the arctic fox was driven by a reduction in home range size in the goose colony. As a result, the net interaction from the presence of geese was negative. Our results also showed that this interaction could lead to local exclusion of sandpipers over a range of adult sandpiper annual survival observed in the wild, which is coherent with previous observations of their co-distribution. Our approach takes into account diverse proximate mechanisms underpinning interaction strengths in a multi-prey system and generates novel insights on some of the predator behavioral responses that may influence prey coexistence (and the lack of) in vertebrate communities.

## 2 Introduction

Understanding how and to what extent biotic interactions influence species occurrence is a major challenge because of the myriad of ways species interact in natural communities (Godsoe et al., 2017). Indirect biotic interactions are especially hard to tackle because they arise through chains of direct interactions (Cazelles et al., 2016). In theory, negative indirect interactions between species that share a common predator (hereafter predator-mediated interactions) may alter community composition by excluding species that are more vulnerable to predation. Although such indirect interactions are likely widespread (Holt and Bonsall, 2017), they are difficult to quantify in complex natural communities (e.g., Iles et al. 2013; Schmidt and Ostfeld 2008; Suraci et al. 2014; Wilson et al. 2022).

Predator-mediated interactions can be quantified according to the change in the number of prey acquired per predator per unit of time (the functional response) and to the change in the number of predators (the numerical response) as a function of prey density. The net effect of the indirect interaction on a given prey species can be either null, negative (e.g., apparent competition) or positive (e.g., apparent mutualism) depending on the relative strength of the predator functional and numerical response (Holt and Bonsall, 2017). For instance, increasing the abundance of a prey *i* should theoretically release predation on prey *k* via the predator acquisition rate (Murdoch, 1969; Murdoch and Oaten, 1975). Alternatively or additionally, increasing the density of a prey *i* could increase the density of predators, and consequently increase predation rate on prey *k* (Holt, 1977; Holt and Lawton, 1994). The balance between such opposing indirect effects has been well studied theoretically (Abrams and Matsuda, 1996) but theoretical predictions have rarely been tested in natural communities. This is in part due to difficulties in obtaining empirically, based multi-species functional response models (Abrams, 2022; DeLong, 2021) and in measuring the relative effects of the predator functional and numerical responses on the net interaction strength. Process-based mechanistic models (hereafter referred to as mechanistic models) can allow us to disentangle the relative strength of the functional and numerical responses of predators, and ultimately improve our ability to accurately quantify the strength of the net indirect interactions in ecological communities (Beardsell et al., 2022; Wootton et al., 2021).

An increase in prey densities may result in higher predator density through behavioral or demographic processes. In most predator-prey models, the numerical response of a predator is incorporated through reproduction and survival parameters (Abrams, 2022; Courchamp et al., 2003; Rosenzweig and MacArthur, 1963; Serrouya et al., 2015). Although a change in prey density is likely to influence the predator density via reproduction or survival, changes in predator behavior can also lead to marked changes in predator density. For instance, an increase in prey density modifies the costs and benefits of movements and competitive interactions, with direct effects on both home range size and local density (Loveridge et al., 2009; Payne et al., 2022). Although this idea is intuitive, the link between predator home range size and predator density is rarely explicitly incorporated in predator-multi-prey models. Yet, this is important to understand the mechanistic processes and model the net effect of predator-mediated interactions in natural communities.

Our objectives were twofold. First, we built a multi-prey mechanistic model of predation by breaking down every step of the predation process to assess whether the presence of a prey species *i* affects acquisition rates of a prey species *k* by a shared predator. We then calculated the resulting predation rates by considering changes in predator density associated with an adjustment in predator behavior (reduction in home range size) induced by the presence of prey *i*. Second, we used a population matrix model to evaluate whether changes in predation rates caused by the presence of prey *i* can indirectly generate the local exclusion of prey *k*. This was illustrated in an arctic vertebrate community composed of a generalist predator, the arctic fox (*Vulpes lagopus*), feeding primarily during the summer on small cyclic mammals and eggs of various tundra bird species, including colonial nesting geese (prey *i*) and sandpipers (prey *k*).

The focal High Arctic community is characterized by high-amplitude fluctuations of lemming populations (with peaks occurring every 3–4 years), and by the presence of a large breeding colony of Snow Geese (Gauthier et al., 2013). In this community, the occurrence probability of nesting shorebirds decreases when colonial nesting geese are present, and shorebird nest predation risk (measured with artificial nests) is higher at high goose nest densities (Duchesne et al., 2021; Lamarre et al., 2017; McKinnon et al., 2013). Although the time required to handle goose eggs can reduce the time available for searching other prey like sandpiper nests, we predicted that this positive effect can be outweighed by an increase in predator density in the goose colony associated with a reduction in fox home range size (Fig. 1). We expected that the resulting predation rates in the presence of the goose colony can be high enough to induce sandpiper local exclusion (without sandpiper immigration). The originality of this study lies in our ability to identify dominant mechanisms affecting prey coexistence (and the lack of) in a natural vertebrate community using models parameterized from a combination of behavioral, demographic, and experimental data acquired over 25 years (Beardsell et al., 2021; Gauthier et al., 2013; Weiser et al., 2020).

**Figure 1:**
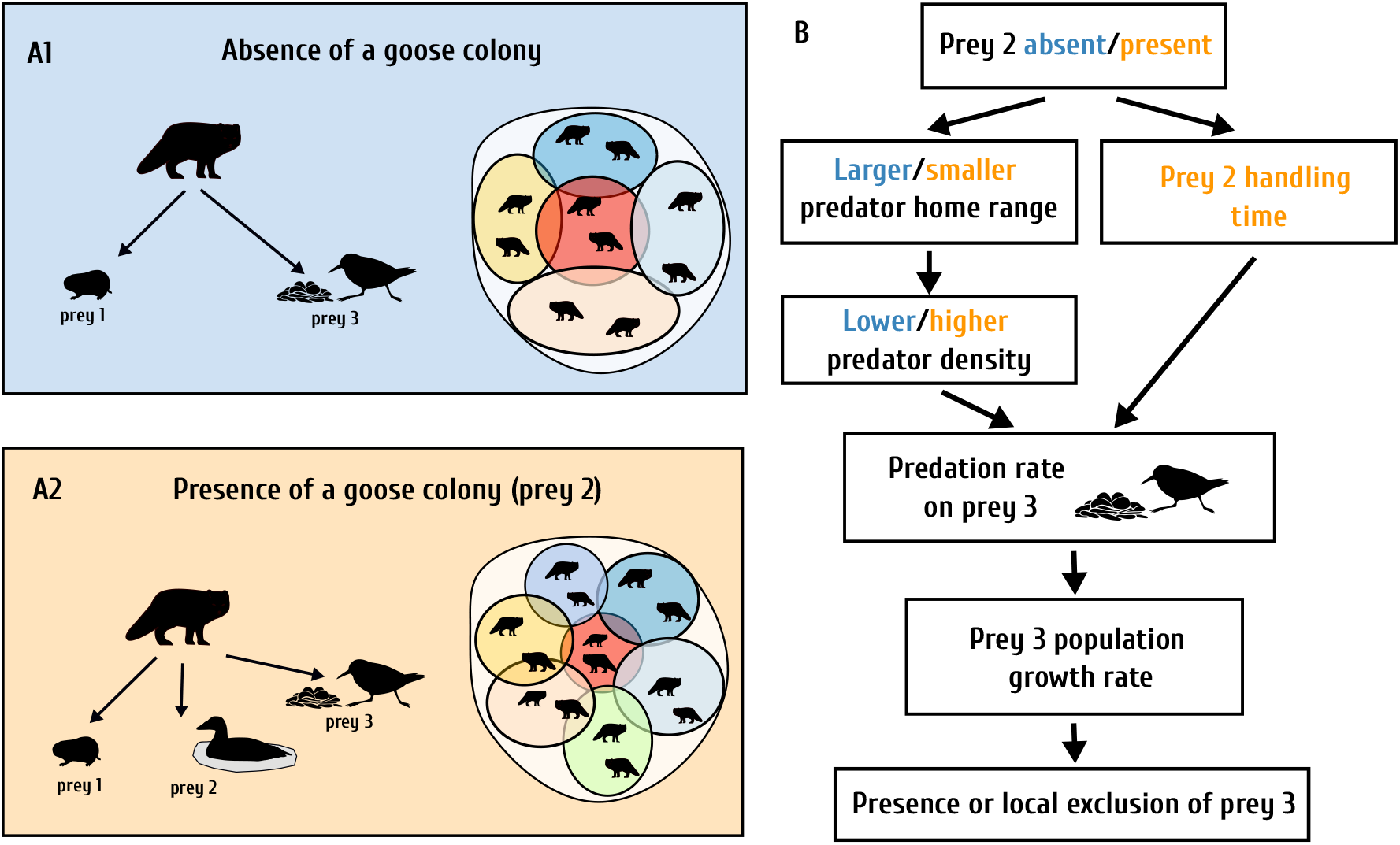
**(A)** Diagrams of simplified Arctic food webs and of fox home range size showing direct links between a predator (Arctic fox), prey 1 (lemmings), prey 2 (goose eggs), and prey 3 (sandpiper eggs) in the absence (**A1**) and presence of a goose colony (**A2**). (**B**) Schematic representation of hypothesized mechanisms underlying the indirect interaction of prey 2 (goose eggs) on prey 3 (sandpiper eggs) through a shared predator (arctic fox). Although the time required to handle goose eggs can reduce the time available searching for sandpiper nests, we predicted that this positive effect can be outweighed by an increase in predator density in the goose colony associated with a reduction in fox home range size.

## 3 Methods

### 3.1 Study area and species

The mechanistic model of predation was built using detailed empirical data from a long-term ecological study on Bylot Island, Nunavut, Canada (73° N; 80° W). The study area (~ 500 km^2^) encompasses a greater snow geese colony (*Anser caerulescens atlanticus*) of ~ 20 000 pairs, which is concentrated in an area of 50-70 km^2^ (McKinnon et al., 2014). The location of the goose colony centroid is relatively stable across years (Duchesne et al., 2021). Two cyclic species of small mammals are present: the brown (*Lemmus trimucronatus*) and collared (*Dicrostonyx groenlandicus*) lemmings (Gauthier et al., 2013). The most common ground-nesting sandpipers found in the study area are the Baird’s (*Calidris bairdii*) and White-rumped (*Calidris fuscicollis*) sandpipers. The arctic fox is an active-searching predator (Poulin et al., 2021) and the main predator of goose and sandpiper eggs (Bêty et al., 2002; McKinnon and Bêty, 2009). In the study area, arctic foxes use the same home range during the summer, and the degree of overlap is generally low in the study population (Clermont et al., 2021).

### 3.2 Multi-prey mechanistic model of predation

We built on a mechanistic model of arctic fox functional response to lemming and sandpiper nests developed within the same study area (Beardsell et al., 2022). We incorporated goose nests into this model based on a mechanistic model previously developed for the fox-goose dyad (Beardsell et al., 2021). The model was derived by breaking down fox predation into a maximum of 6 steps: (1) search, (2) prey detection, (3) attack decision, (4) pursuit, (5) capture and (6) manipulation. Each step was adapted to each prey species according to their anti-predator behavior and the fox hunting behavior (Beardsell et al., 2021, 2022). Figure 2 provides an overview of the multi-prey mechanistic model (prey 1 is lemmings, prey 2 is goose nests and prey 3 is sandpiper nests). Detailed equations of the predation model can be found in Appendix S2 and associated parameter values in Table 1. For details on the construction of the model and the extraction of parameter values see Beardsell et al. (2021, 2022).

**Figure 2:**
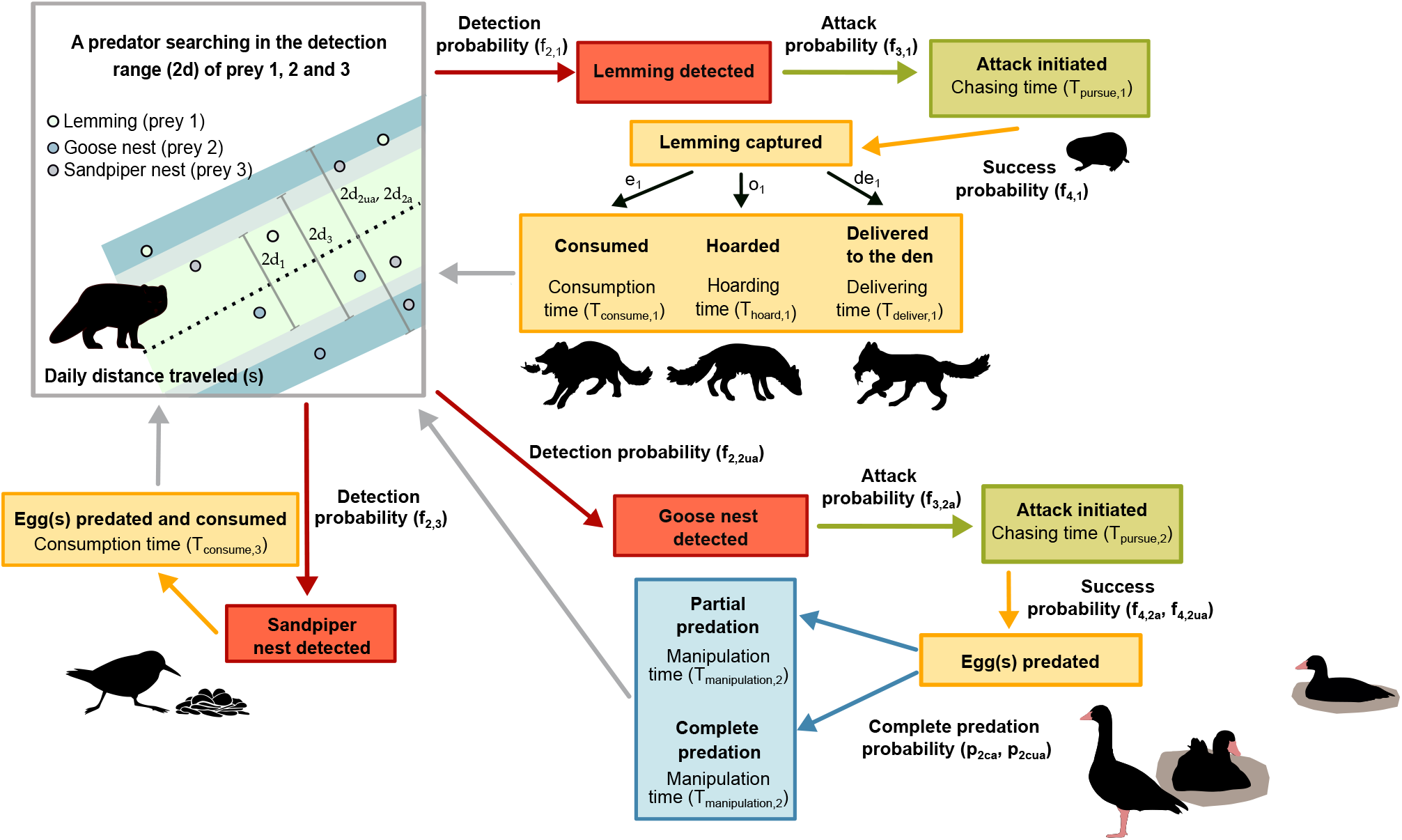
Conceptual multi-prey mechanistic model of arctic fox functional response to density of lemmings (prey 1), goose eggs (prey 2) and sandpiper eggs (prey 3). Each box represents one or more components of predation (search, prey detection, attack decision, pursuit, capture and manipulation). Arrows represent the probability that the predator reaches the next component. When there is no parameter near the arrow, the probability of reaching the next component is 1. As incubating geese can actively protect their nests from arctic foxes, their presence at the nest strongly influences fox foraging behavior. Thus, most parameter values were estimated separately for goose nests that were attended and unattended (indicated by two symbols near the arrows).

We calculated sandpiper nest predation rate in the presence of a goose colony over a 50 km^2^ area (*A*; which is the average core area of the goose colony (Duchesne et al., 2021)) and an equivalent area where geese are absent. The number of sandpiper nests predated per day within *A* is given by the product of predator acquisition rate (namely the functional response) and the number of foxes present in *A*. The equation describing predation rate on sandpiper nests (*P_3_* (*N*_1_, *N*_2_, *N*_3_); nests day^-1^ within *A*) is:

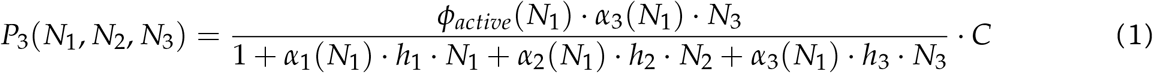

Where *ϕ_active_* is the proportion of time the predator spent active in a day, N the density of each prey (ind/km^2^), *α* the capture efficiency (km^2^/day), and *h* the handling time per prey item (day/per prey item). For instance, capture efficiency of a lemming (prey 1) is expressed by the product of the daily distance traveled by the predator (*s*; km^2^/day), the reaction distance (*d*_1_; km), the detection probability (*f*_2,1_), the attack probability (*f*_3,1_) and the success probability of an attack (*f*_4,1_). The capture efficiency equations for other prey can be found in the Appendix S2 and associated parameter values in Table 1. Since we have evidence that the values of *ϕ_active_* and *s* depend on lemming density (Model C in Beardsell et al. 2022), the value of *ϕ_active_* and *α* are expressed as a function of lemming density. The predation rate on sandpiper nests in absence of a goose colony is obtained by setting the density of geese (*N*_2_) to 0 in equation 1.

We estimated the number of predators (C) within *A* as follows:

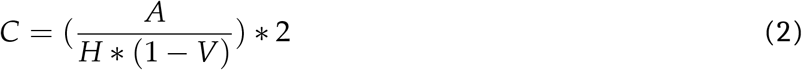

Where *H* is the average home range size (km^2^), and *V* is the average proportion of overlap between adjacent home ranges. Based on high-frequency gps-data, the average overlap between adjacent home ranges is 0.18 on Bylot Island (Clermont et al., 2021). We estimated summer home range size of arctic foxes using telemetry data (Argos) of 113 foxes from 2008 to 2016 on Bylot Island. Foxes were captured and equipped with Argos radio collars as described in Tarroux et al. (2010), providing a location every 1-2 days. We estimated the area of the 95% home range contour for each individual-year between May-October using the autocorrelation-informed home range estimation workflow described in Fleming et al. (2015), and implemented in *ctmm* R package (Calabrese et al. 2016, Dulude et al. in prep.). Home range size averages 10.8 km^2^ (*n* = 56 home ranges) and 18.2 km^2^ (*n* = 57) in presence and in absence of the goose colony respectively (Fig. 3A). As fox pair members share a home range (Clermont et al., 2021), we assumed that two foxes were foraging per home range.

**Figure 3:**
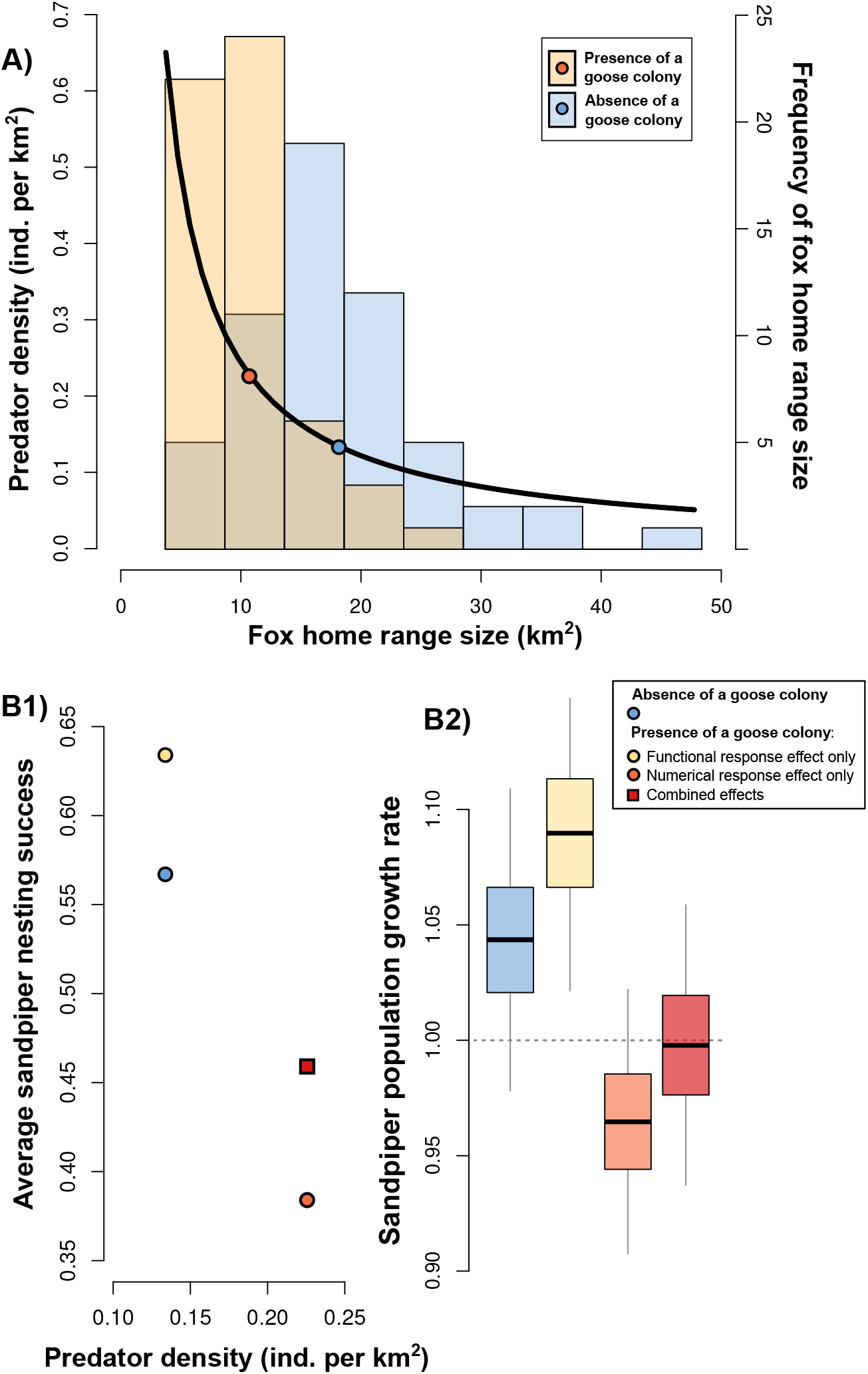
(**A**) Relationship between fox density and summer home range size of arctic foxes derived from equation 2, and histograms of home range size (estimated from telemetry data; n = 113) in presence and in absence of a goose colony. Points indicate average home range size in presence and in absence of a goose colony. (**B1**) Relationship between average nesting success of sandpipers (calculated for a 13 year period covering different lemming densities; see methods) and fox density in the absence and presence of geese (functional response effect only, numerical response effect only and combined effects). (**B2**) The effects of goose presence (functional response effect only, numerical response effect only and the combined effects) on local growth rate (*λ*) of sandpipers calculated with the population matrix model. The center line shows the median, the boxes show the interquartile range, and the lines show the minimum and maximum values.

We estimated annual success of sandpiper nests (prey 3) in the presence and in the absence of a goose colony from daily predation rates (Eq.1) using a set of differential equations. These equations allowed us to calculate the predator acquisition rate over the sandpiper nesting period (i.e., the average duration between the laying date and hatching date; 24 days). We assumed that the bird nesting period is synchronized and that predated nests are not replaced. Thus, the density of goose and sandpiper nests decreases each day, as expressed by Equation 3. The number of nests predated after 24 days is then divided by the maximum number of nests present at the first day of nest initiation (*Q*; calculated by the product of the density of *N*_3_ on day 1 and *A*), giving us an estimate of annual predation rate (annual nesting success is obtained by subtracting the predation rate by 1). Sandpiper nests predation rate in the presence of a goose colony is given by *P*_3_(*N*_1_, *N*_2_, *N*_3_) * *C*. The rate of change in sandpiper nest density (*N*_3_) is expressed as follows:

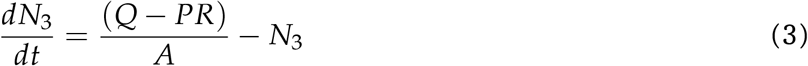

Where *A* is the average core area of the goose colony (50 km ^2^). Equivalent set of equations for predation rate in the absence of the goose colony can be obtained by setting the density of geese (*N*_2_) to 0. We also estimated annual success of goose nests (prey 2) by substituting all 3 for 2 and vice versa in equations **??** and 3, and over a period 28 days (instead of 24 days to correspond to the average duration between the laying date and hatching date).

### 3.3 Sensitivity analysis

We quantified the relative influence of model parameter values on the estimation of sandpiper annual nesting success by using the Latin hypercube sampling technique (an efficient implementation of the Monte Carlo methods; Marino et al. 2008). This analysis allowed us to investigate the uncertainty in the model output generated by the uncertainty and variability in parameter inputs. Each parameter was represented by a probability distribution (uniform or normal truncated) based on the distribution of empirical data. For some parameters, the biological information was limited, so we assigned a uniform distribution allowing for a large range bounded by minimum and maximum values. Latin hypercube sampling was then applied to each distribution (N = 1,000 iterations). For simplicity, the sensitivity analysis was conducted on the predation model without density-dependence in parameters *s* and *ϕ_active_* and all prey were set at intermediate densities (*N*_1_ = 350 individuals/km^2^, *N*_2_ = 255 nests/km^2^, *N*_3_ = 3.1 nests/km ^2^).

### 3.4 Simulations

We calculated average nesting success of sandpipers for average arctic fox home range sizes, as well as for the whole range of home range sizes observed in the presence and absence of a goose colony. Since lemming densities fluctuate with high amplitude between years, we computed the average sandpiper nesting success over the 13-year time series of lemming densities on Bylot Island (Appendix 1 Fig. S1). As expected due to Jensen’s inequality (Ruel and Ayres, 1999), inclusion of interannual variability in lemming density (from 2 to 648 ind./km^2^) results in a 7% decrease in average nesting success of sandpipers relative to a constant average lemming density (i.e., 204 ind/km^2^). We set the average density of goose nests within fox home ranges at 255 nests per km^2^. We derived this estimate from an exhaustive count of all goose nests present within the colony (an area of 56 km^2^ in 2019; methods in Grenier-Potvin et al. 2021). The year 2019 falls within the long-term average of goose nest density measured in an intensive monitoring area (0.5 km^2^) in the core of the colony from 1989-2019 (Gauthier et al., 2019). We conducted all models and simulations in *R* v. 4.0.4 (R Core Team, 2021).

### 3.5 Sandpiper population model

We evaluated if changes in nest predation rates caused by the presence of a goose colony can indirectly generate local exclusion of sandpipers. We used a population matrix model to link estimated nesting success to sandpiper population growth rate. Since most demographic parameters for white-rumped sandpiper and/or Baird’s sandpiper are poorly documented on Bylot Island, we build upon a projection matrix model developed by Weiser et al. (2020) for the semipalmated sandpiper (*Calidris pusilla;* hereafter sandpiper), a tundra nesting species for which the demographic parameters are relatively well documented across the North American Arctic (Appendix S3; Fig. S1). We calculated growth rate (*λ*) using the mean values of each vital rate while varying average nesting success values (*NS_ini_* and *NS_renest_*; see Table S1 in Appendix S3). Given the strong influence of annual adult survival on *λ* and since empirical data of nesting probability are virtually absent (Weiser et al., 2020), we conducted simulations for various values of annual adult sandpiper survival (0.76 ± 5%) and nesting probability (from 0.8 to 1). We used the *popbio* package v. 2.7 (Stubben and Milligan, 2007) in *R* (R Core Team, 2021) to calculate *λ*. Details regarding the matrix model are available in Appendix S3.

**Table 1:** Symbol definition and parameter values used in the multi-prey mechanistic model of fox predation as a function of the density of lemmings (prey 1), goose nests (prey 2) and sandpiper nests (prey 3). Parameter values were estimated from a combination of high-frequency GPS and accelerometry data (23 summer foxes, 2018-2019), ARGOS telemetry data (113 summer-foxes), behavioral observations in the field (*n* = 124 hours, 1996-2019) and camera traps deployed at nests (2006-2016). Most details regarding the estimation of parameter values can be found in Beardsell et al. (2021). Parameters related to lemming manipulation times and the fox activity budget can be found in Beardsell et al. (2022).

| Parameter name | Symbol | Value(s) | Unit |
| --- | --- | --- | --- |
| **Arctic Fox** |  |  |  |
| Home range size | $H$ | 3.7-48.4 | km<sup>2</sup> |
| Average proportion of overlap between adjacent home ranges | $V$ | 0.18 | |
| Proportion of time the predator spent active in a day | $\phi_{active}$ | 0.5 | - |
| Daily distance traveled by the predator (when $\phi_{active} = 0.5$) | $s$ | 41 | km day<sup>-1</sup> |
| **Lemmings** |  |  |  |
| Lemming density | $N_1$ | 0-700 | ind. km<sup>-2</sup> |
| Maximum reaction distance | $d_1$ | 0.0075 | km |
| Average detection and attack probability within the reaction distance | $f_{2,1} * f_{3,1}$ | 0.15 | - |
| Success probability | $f_{4,1}$ | 0.51 | - |
| Chasing time | $T_{pursue,1}$ | $1.0 \times 10^{-3}$ | day ind.<sup>-1</sup> |
| Consumption time | $T_{consume,1}$ | $3.8 \times 10^{-4}$ | day ind.<sup>-1</sup> |
| Consumption probability | $e_1$ | 0.48 | - |
| Hoarding time | $T_{hoard,1}$ | $4.9 \times 10^{-4}$ | day ind.<sup>-1</sup> |
| Hoarding probability | $o_1$ | 0.32 | - |
| Delivering time | $T_{deliver,1}$ | $3.9 \times 10^{-3}$ | day ind.<sup>-1</sup> |
| Delivering probability | $de_1$ | 0.20 | - |
| **Goose nests** |  |  |  |
| Goose nest density | $N_2$ | 255 | nests km<sup>-2</sup> |
| Nest unattendance probability | $w$ | 0.021 | - |
| Chasing time | $T_{pursue,2}$ | $8.3 \times 10^{-4}$ | day nest<sup>-1</sup> |
| Manipulation time (includes consumption and hoarding time) | $T_{manipulation,2}$ | $5.8 \times 10^{-3}$ | day nest<sup>-1</sup> |
| **Goose attended nests** |  |  |  |
| Maximum reaction distance | $d_{2a}$ | 0.033 | km |
| Average attack probability within the reaction distance | $f_{3,2a}$ | 0.05 | - |
| Success probability | $f_{4,2a}$ | 0.098 | - |
| Complete predation probability | $p_{2ca}$ | 0.47 | - |
| **Goose unattended nests** |  |  |  |
| Maximum reaction distance | $d_{2ua}$ | 0.11 | km |
| Average detection probability within the reaction distance | $f_{2,2ua}$ | 0.37 | - |
| Success probability | $f_{4,2ua}$ | 0.93 | - |
| Complete predation probability | $p_{2cua}$ | 0.69 | - |
| **Sandpiper nests** |  |  |  |
| Sandpiper nest density | $N_3$ | 3.1 | nests km<sup>-2</sup> |
| Maximum reaction distance | $d_3$ | 0.085 | km |
| Average detection probability within the reaction distance | $f_{2,3}$ | 0.029 | - |
| Consumption time | $T_{consume,3}$ | $2.8 \times 10^{-3}$ | day nest<sup>-1</sup> |

## 4 Results

Summer home range size of arctic foxes varied from 3.7 to 48.4 km^2^ in the study area (Fig. 3A). The average home range size was smaller whithin the colony (18.2 km^2^ outside vs 10.8 km^2^ within the goose colony), and hence the estimated fox density was on average 1.7 times higher in the goose colony (Fig. 3A). The estimated goose nesting success was 76%, which is consistent with the average success estimated from intensive annual goose nest monitoring in the colony (68% between 1991 and 2015; Reséndiz-Infante et al. 2020). In absence of nesting geese, the estimated nesting success of sandpiper was 56% (Fig. 3B). This is also consistent with the average nesting success observed in a monitoring area located ~30 km away from the goose colony on Bylot Island (50% ± 0.08 (SE) between 2005 and 2019; Beardsell et al. 2022). There is no observation of of sandpiper nesting success in the goose colony because sandpiper nest density is too low (Lamarre et al., 2017).

Sensitivity analysis indicated that four parameters had a significant effect on annual sandpiper nesting success (Appendix S1; Fig. S2). A change in the value of these four parameters by 50% generated changes in sandpiper annual nesting success by 24%, 15%, 12%, and 11% respectively for predator home range size, predator speed, proportion of time spent active by the predator, and detection probability of sandpiper nests (Appendix S1; Fig. S3). Neither speed nor activity level was correlated to fox home range sizes based on high-frequency GPS and accelerometry data (Appendix Fig. S5). Although some parameters directly related to goose nest predation had a statistically significant influence on sandpiper nesting success (Appendix S1; Fig. S2), their biological effects were limited as indicated by the low correlation coefficient (<0.21) of the relationship (Appendix S1; Fig. S3). Predator home range size was thus the most influential parameter in the model.

We evaluated the net effect of colonial geese on the average sandpiper nesting success. We first computed nesting success of sandpipers from the multi-prey mechanistic models over the range of fox densities (home range sizes) observed in the study area (Appendix S1 Fig.S4). For a given arctic fox density, the presence of nesting geese increased the estimated sandpiper nesting success by 7% (functional response effect only: Fig. 3B1; see also Appendix S1 Fig.S4). This release of predation pressure was the result of time constraints related to goose egg handling (including chasing, hoarding, consumption), which reduced the time available for searching other prey like sandpiper nest. On the other hand, when considering only the increase in fox density caused by the presence of colonial geese (from 0.13 ind./km^2^ to 0.22 ind./km^2^; Fig. 3A), the estimated sandpiper nesting success decreased by 18% (numerical response effect only: Fig. 3B1). The negative effect mediated by arctic fox home range size adjustment thus outweighed the predation release due to goose egg handling time, resulting in an 11% decrease in average sandpiper success in the goose colony overall (see combined effects in Fig. 3B1).

We investigated the net effect of the goose colony on sandpipers demography. Population growth rate (*λ*) derived from the sandpiper matrix population model indicated that changes in sandpiper nesting success caused by the presence of colonial geese can affect local sandpiper population dynamics (Fig. 3B2). While the predation release on sandpiper nests generated by the goose egg handling time could increase *λ* by 3% (functional response effect only), the reduction in sandpiper nesting success caused by higher density of foxes in the goose colony resulted in a 7% decrease of *λ* (numerical response effect only: Fig. 3B2). The negative effect mediated by the increased predator density thus outweighed the positive effect generated by the functional response. The combinaison of these effects in presence of geese is sufficient to drive sandpiper local exclusion for various combinations of adult sandpiper annual survival and nesting probability whereas in absence of geese *λ* >1 (Fig. 3B2). For an average fox home range size observed in the goose colony on Bylot Island, model outputs indicated that sandpiper adult survival has to reach a minimum of 0.74 for a *λ* >1 without immigration (Appendix S1; Fig. S6).

## 5 Discussion

In this study, we used a mechanistic multi-prey predation model to quantify predator-mediated interaction strength in a natural system. We analyzed the model to quantify the indirect interaction between two prey species (colonial nesting geese and sandpipers) sharing a common predator (the arctic fox). We incorporated predation rates into a population matrix model to evaluate the consequences of predator-mediated interactions on local prey growth rates. Our results showed that the positive effects of the presence of a goose colony on sandpiper nesting success (due to the handling time of goose eggs by the predator) were outweighed by the negative effect of an increase in fox density, associated with a reduction in fox home range size in the goose colony. Thus, the net interaction resulting from the presence of a goose colony on sandpiper nesting success was negative. The strength of the negative net interaction obtained could be sufficient to cause local exclusion of sandpipers for various values of adult sandpiper survival rate observed in the wild. Overall, our results indicate that predator-mediated effects could explain the low occurrence of Arctic-nesting shorebirds in areas of high goose nesting density (Duchesne et al., 2021; Flemming et al., 2016; Lamarre et al., 2017).

The strength of the negative indirect interaction of geese on sandpipers was potentially underestimated due to a combination of factors. First, in addition to reducing home range size, the presence of abundant resources could also increase overlap between fox home ranges (Eide et al., 2004; Lai, 2017). This was not taken into account in our models because adjacent fox home ranges were not systematically monitored. Second, in addition to causing lower egg survival, higher fox density is likely to reduce chick survival. However, empirical data on chick survival is limited since sandpiper chick leave the nest shortly after hatching. Finally, a higher density of avian predators within the goose colony (Lamarre et al., 2017) may also decrease survival rate of sandpiper chicks. These three factors would have amplified the strength of the negative effect of the presence of geese on sandpipers if they had been included in the model.

Along with changes in the predator home range size, additional components of predator behavior are likely to change in the presence of geese and more data is needed to fully explore the possible links between those parameters. Our sensitivity analysis indicated that three parameters have a notable influence on sandpiper nesting success, namely 1) daily distance traveled by the predator, 2) proportion of time spent active by the predator, and 3) sandpiper nest detection probability by the predator. We recognize that further field investigations, such as long-term GPS and accelerometer tracking of predators over a wide range of prey densities, are needed to investigate the effect of prey densities on the value of predator movement parameters. This would be especially important in our study system since changes in these movement parameters are known to be related to lemming density (Beardsell et al., 2022). Regarding the detection probability of a sandpiper nest, there is no evidence that this parameter is affected by the presence of geese. This absence of effect probably reflects that attacking a sandpiper nest provides systematic benefits to foxes and entails very low costs (e.g., risk of injury, handling time; Beardsell et al. 2022).

Predator-mediated interactions in natural systems have been investigated using various approaches, including statistical analyses linking prey occurrence probability with density of other prey (Duchesne et al., 2021; Flemming et al., 2019), and field experiments involving the addition or removal of prey or predator species (Menge, 1995; Spiller and Schoener, 2001). Although these approaches can help identifying the presence of indirect effects, they provide a limited ability to tease apart and infer proximate mechanisms underlying apparent biotic indirect interactions. Moreover, field experiments in natural food webs can be impossible to implement when predator home range size is large (but see Wilson et al. 2022).

Variations in the shape of the functional response can have important ecological consequences for the structure and dynamics of communities by altering the coexistence among prey, and the strength and signs of the interactions among them (Abrams and Cortez, 2015; Abrams et al., 1998; Abrams and Matsuda, 2004; Brose et al., 2006; Coblentz, 2020). However, very few empirically-based multi-species functional responses were developed (Abrams, 2022; DeLong, 2021). The evaluation of functional response using phenomenological models often fails to discriminate between different response shapes (Novak and Stouffer, 2020), which makes it difficult to quantify the strength of predator-mediated interactions in the wild. Although strong empirical foundation on multi-species functional response in natural communities is lacking, they are widely used in predator-prey models (Courchamp et al., 2000, 2003; McLellan et al., 2010; Roemer et al., 2002; Serrouya et al., 2015). To our knowledge, our study provides the first empirically based model that integrates mechanistic multi-species functional responses while also taking into account behavioural processes underlying the numerical response of a generalist predator. This is a major step forward in our ability to accurately quantify the consequences of predation on wild animal community structure and dynamics. By providing important empirical data, the growing number of technologies enabling the remote monitoring of wildlife behavior (e.g., high-frequency GPS, acoustic and heart rate monitors) should facilitate the application of our model in more complex systems (Pagano et al., 2018; Studd et al., 2021; Williams et al., 2014).

Our results show that finer-scale behavioral processes may actually be the main drivers of predator density and prey persistence in the wild. Along with the link between predator home range size and prey availability (Bino et al., 2010; Loveridge et al., 2009; Payne et al., 2022), other processes could be explored such as the presence of predator social or aggressive interactions and predator group hunting. For instance, we might expect overlap between predator home ranges to increase when food resources are low and unpredictable. In arctic foxes, this could occur during years of low lemming density, in absence of a goose colony or when foxes mainly feed on unpredictable prey (e.g., carcasses). Such effects remain to be explored.

As pointed out by Abrams (2022), our understanding of numerical responses is much more limited than functional responses. To date, the numerical response is typically modeled through demographic processes in classical models (see MacArthur-Rosenzweig equations; Rosenzweig and MacArthur 1963). Our approach takes into account diverse proximate mechanisms underpinning interaction strengths in a multi-prey system and generates novel insights on some of the predator behavioral responses that may influence prey coexistence (and the lack of) in vertebrate communities. Overall, this study underlines the need to explicitly investigate the consequences of various behavioral processes underlying predator numerical response.

## 6 Acknowledgments

We are grateful to the many people who helped us with field work over the years, to the Mittimatalik Hunters and Trappers Organization, and to Park Canada staff for their assistance. We also thank Marie-Pier Laplante for the English revision. The research relied on the logistic assistance of the Polar Continental Shelf Program (Natural Resources Canada) and of Sirmilik National Park of Canada. The research was funded by (alphabetical order): Arctic Goose Joint Venture, the Canada Foundation for Innovation, the Canada Research Chairs Program, the Canadian Wildlife Service (Environment Canada), the Fonds de recherche du Québec-Nature et technologies, the International Polar Year program of Indian and Northern Affairs Canada, the Kenneth M Molson Foundation, the Natural Sciences and Engineering Research Council of Canada, the ArcticNet Network of Centers of Excellence, the Northern Scientific Training Program, Polar Knowledge Canada, Université du Québec à Rimouski, Université Laval and the W. Garfield Weston Foundation.

## Appendix S1 Supplementary figures

**Figure S1:**
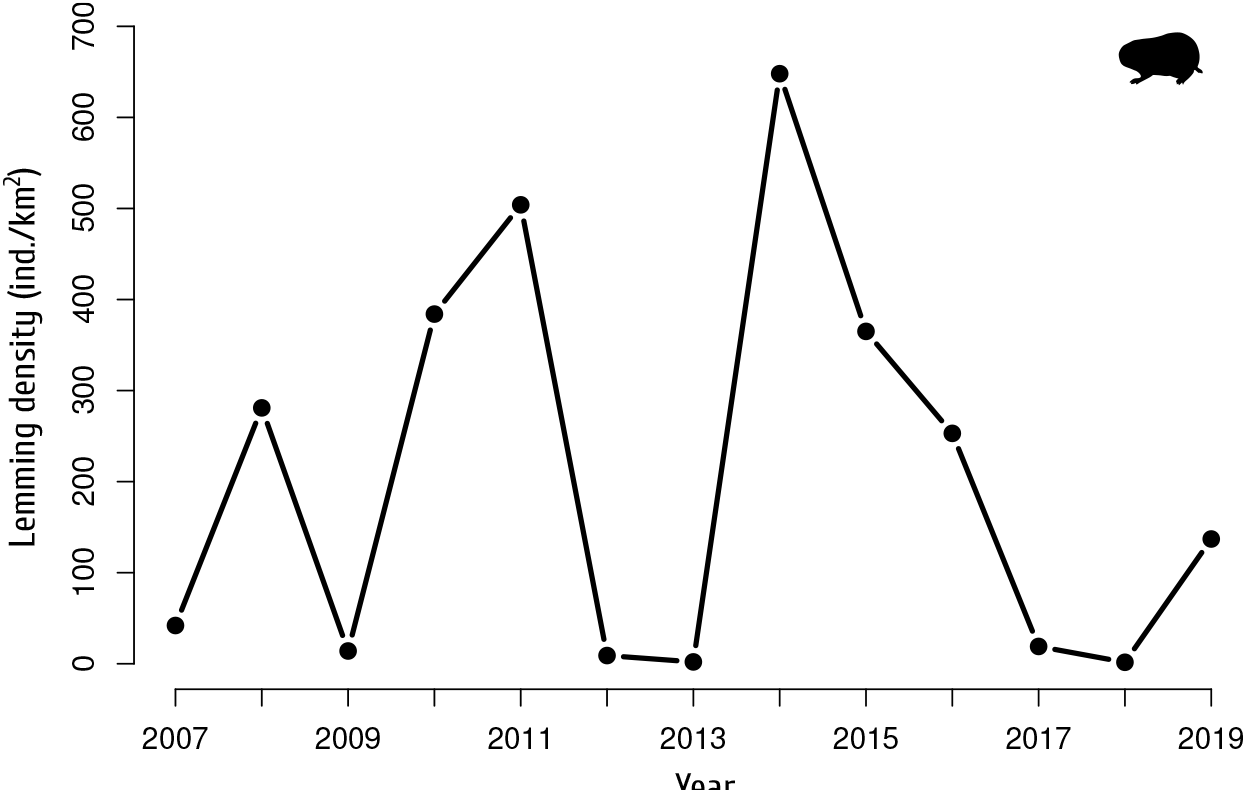
Empirical time series of lemming density on Bylot Island from 2007 to 2019 measured by live-trapping (see methods in Fauteux et al. 2018). The density of brown and collared lemmings was summed.

**Figure S2:**
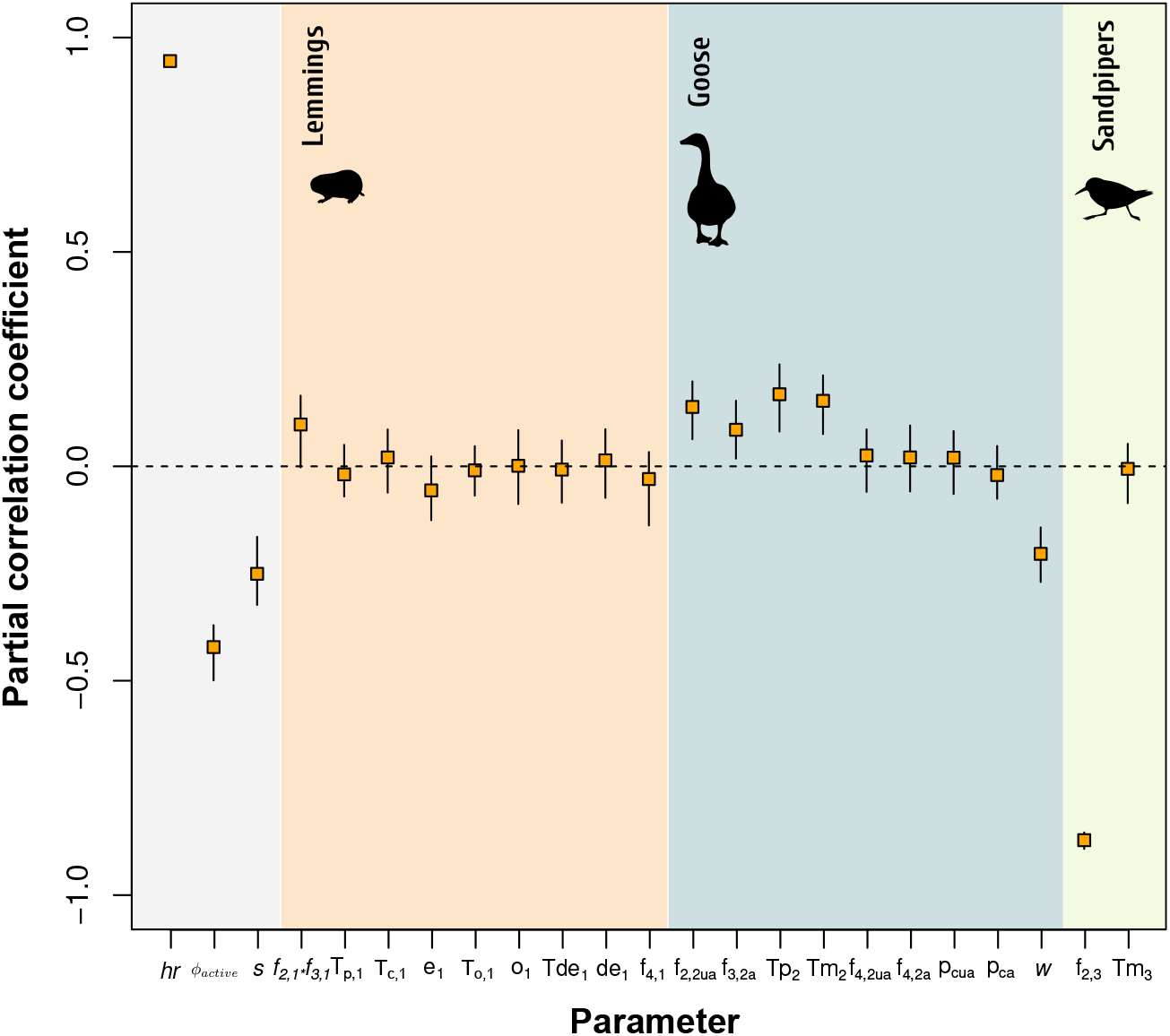
Partial correlation coefficient between the values of each parameter and annual sandpiper nest success. The predation model used in the simulation includes the presence of a goose colony (Eq. 1). The bars are 95% confidence intervals, generated by bootstrapping 40 times (n = 1000 simulations). See Table 1 for a description of each parameter.

**Figure S3:**
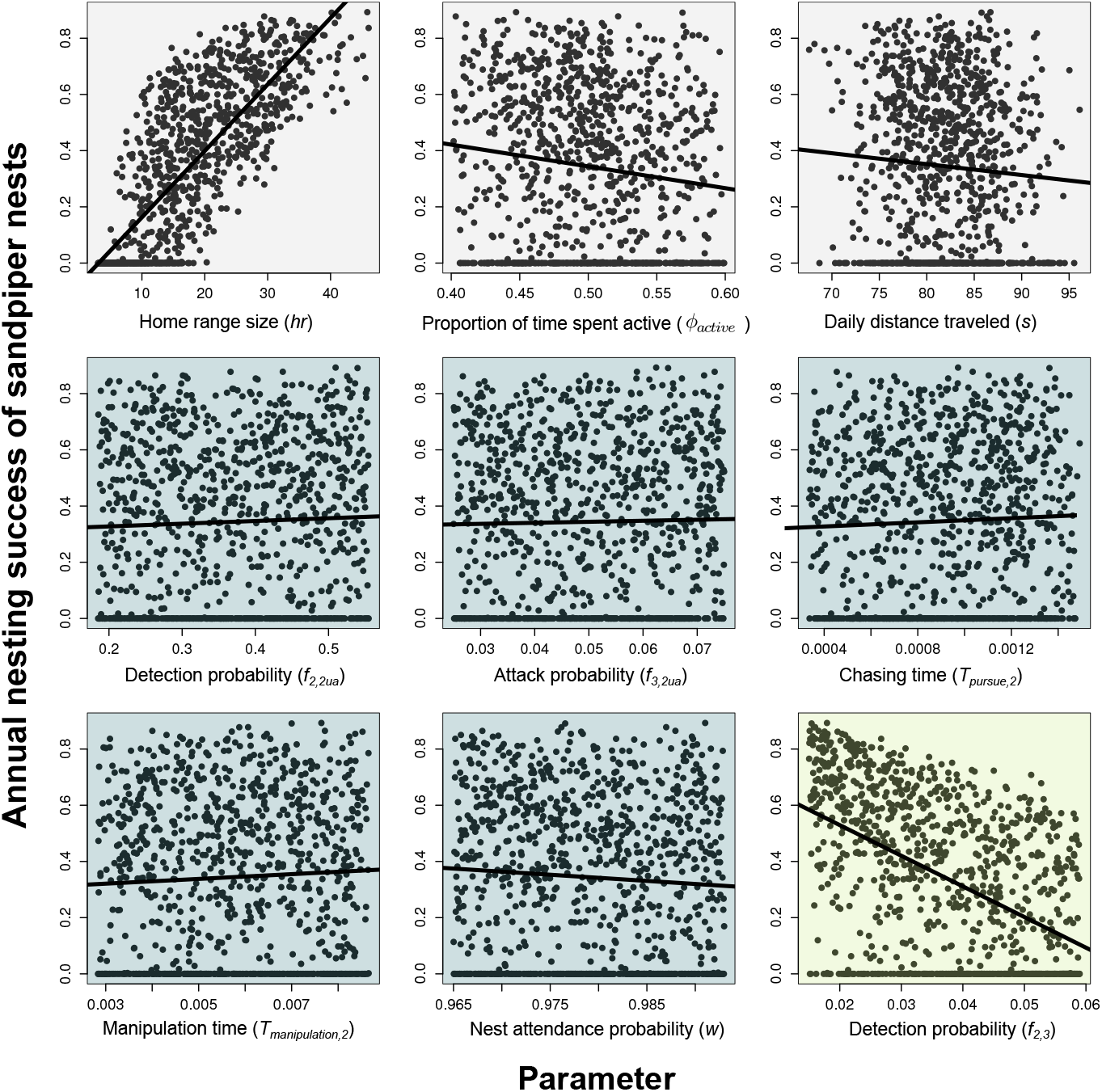
Scatterplots linking the value of input parameters on the annual nesting success of sandpiper nests. Parameters with a correlation coefficient in Fig. S2 significantly different from 0 are represented. See Table 1 for a description of each parameter.

**Figure S4:**
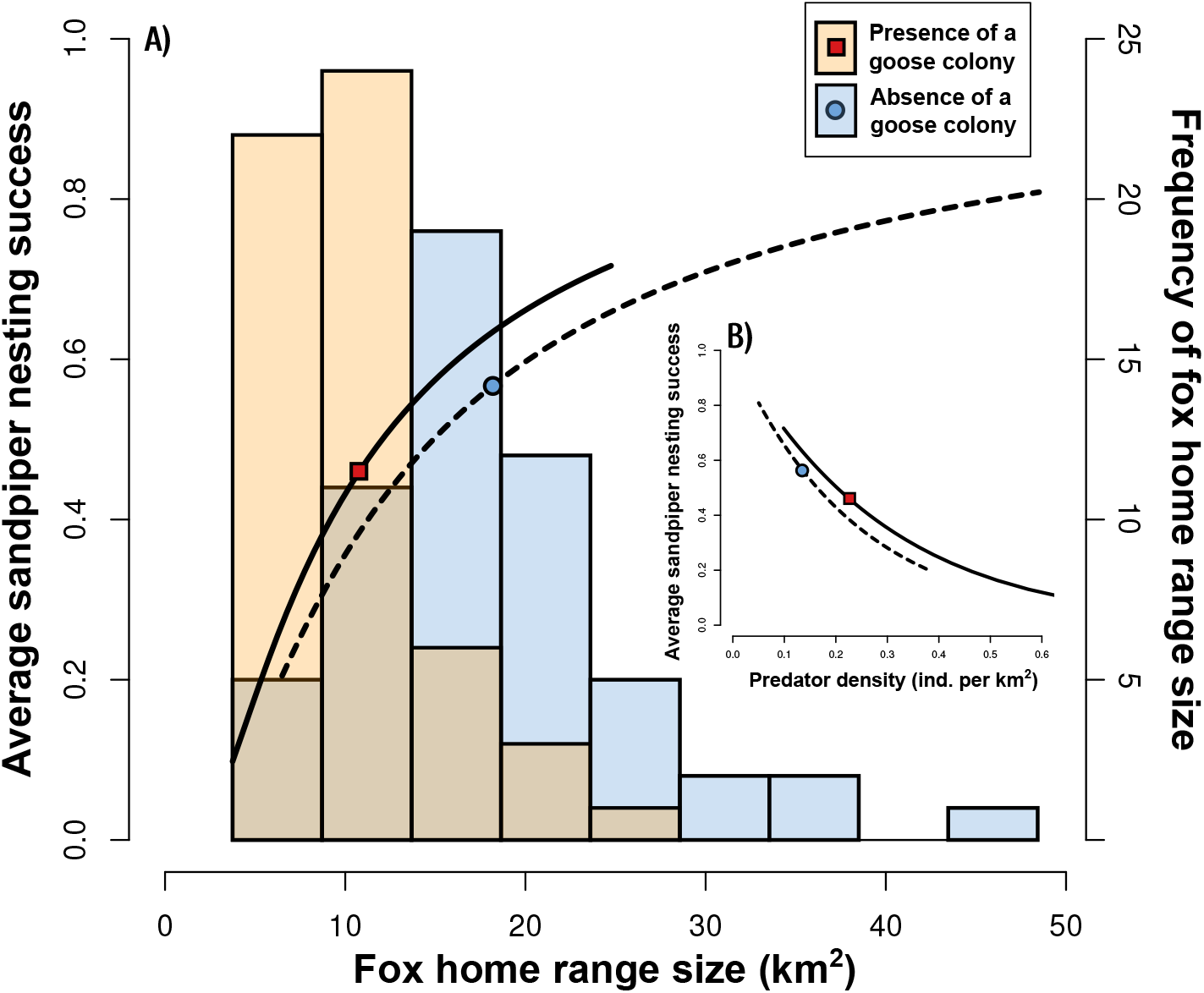
**A)** Histograms of summer home range size of arctic foxes (estimated with telemetry data (*n* = 113)) with and without the presence of a goose colony. Lines indicate the relationship between empirical arctic fox home range size and sandpiper nesting success averaged over 13 years (covering different lemming densities) in the presence (solid line) and absence (dashed line) of a goose colony. The point and square indicate the average nesting success for average home range sizes. **B)** The inset shows the relationship between arctic fox density (calculated from eq. 2) and sandpiper nesting success averaged over 13 years in the presence (solid line) and absence (dashed line) of a goose colony.

**Figure S5:**
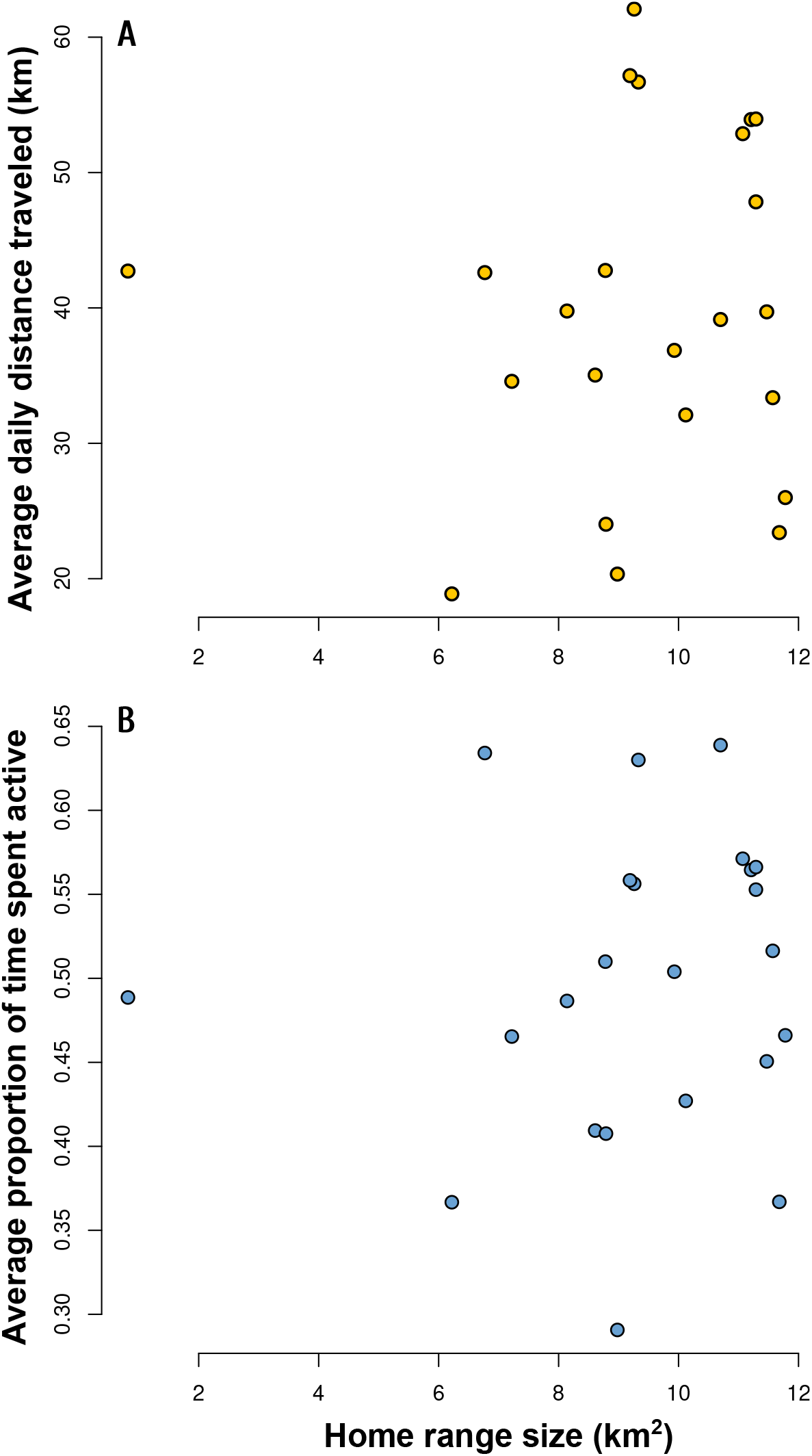
**A)** Relationship between the average daily distance traveled by the arctic fox during the bird incubation period (from June 10 to July 14) and the home range size. **B)** Relationship between the average proportion of time spent active by the arctic fox during the birds incubation period and the home range size. Dots in **A)** and **B)** represent empirical home range size, average daily distance traveled and average proportion of time spent active by the arctic fox calculated from high-frequency GPS-data and accelerometry data from 23 foxes monitored during the summer (8 foxes in 2018 and 15 in 2019).

**Figure S6:**
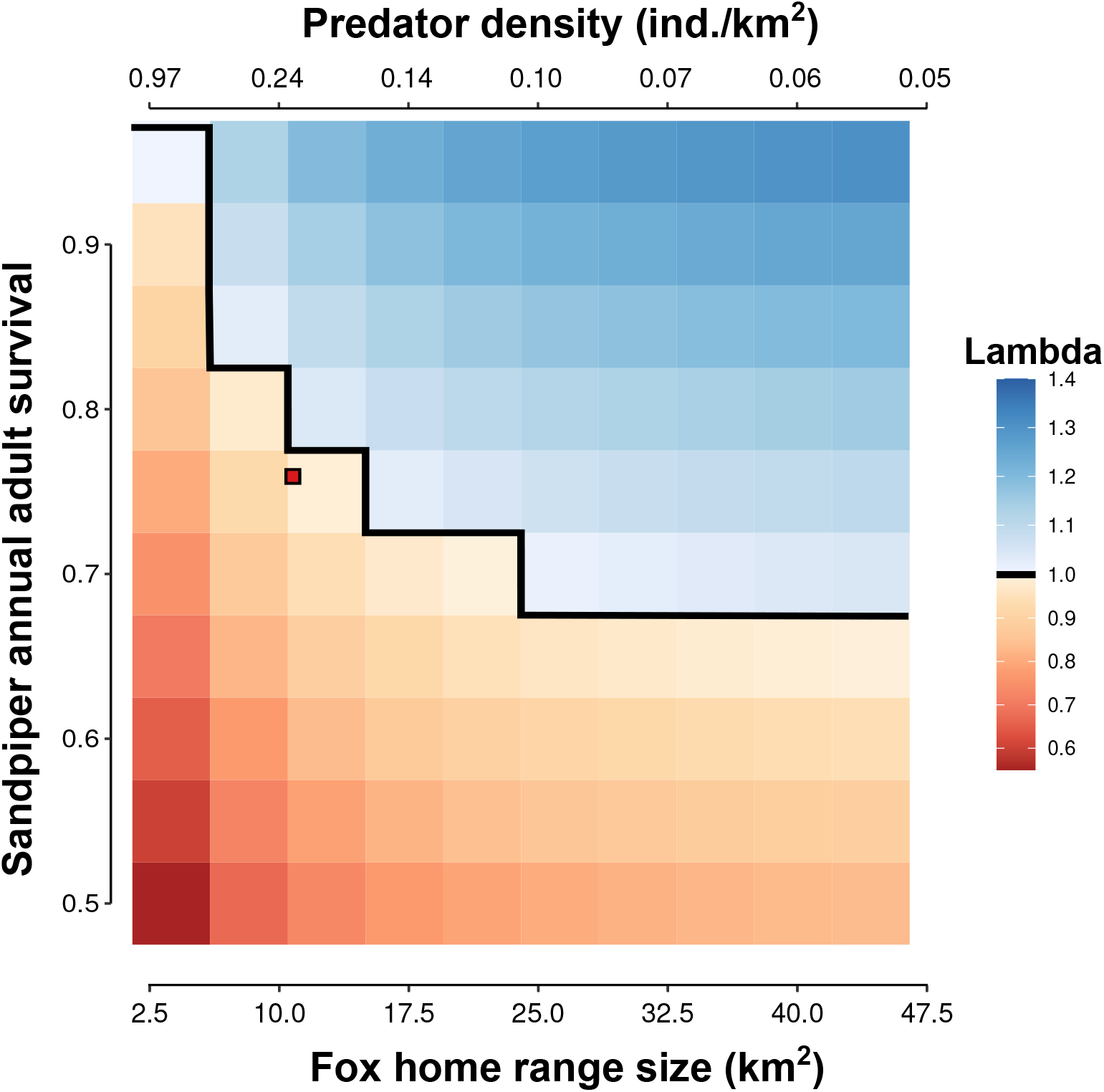
Predicted local growth rate from the population model of semipalmated sandpiper for various combinations of adult survival and fox home range size. The red square is the average home range size observed in the goose colony on Bylot Island along with the average adult survival estimated in semipalmated sandpipers in North America (Weiser et al., 2018). The nesting probability is set to 0.8.

## Appendix S2 Equations of the multi-prey mechanistic model of functional response

This appendix presents mainly the equations. See Beardsell et al. (2021) and Beardsell et al. (2022) for the full details.

### 7.1 Acquisition rate of lemmings by arctic foxes (prey 1)

The number of lemmings captured per fox per day (the acquisition rate) is expressed as:

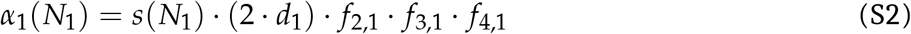

Where capture efficiency (km^2^/day) is defined as:

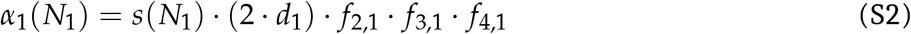

and handling time (day/per lemming) is defined as:

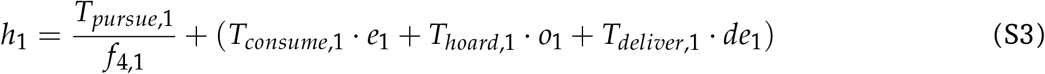

### 7.2 Acquisition rate of goose nests by arctic foxes (prey 2)

Geese can actively protect their nests from arctic foxes, and their presence at the nest strongly influences fox foraging behavior (Bêty et al., 2002). Thus, the model was divided into two components. A first component models the rate of acquisition of goose nests when the female is incubating or when a protective adult is <10 m from the nest (attended nest). A second component models the rate of goose nest acquisition during incubation recesses when both adults are >10 m from the nest (unattended nest). See Beardsell et al. 2021 for more details on the goose model. The total number of goose nests acquired per fox per day (the acquisition rate) is expressed as:

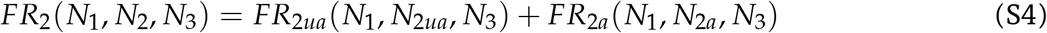

#### 7.2.1 Acquisition rate of unattended goose nests by arctic foxes (prey 2)

The number of unattended nests acquired is expressed as:

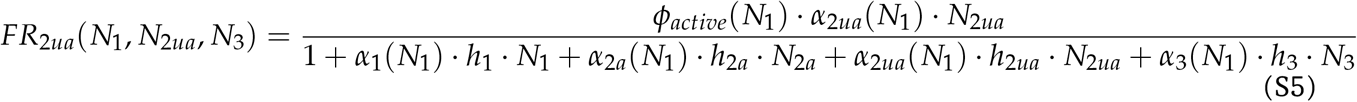

where the density of unattended nests is the product of goose nest density and nest attendance probability (*w*):

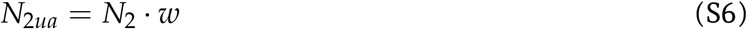

capture efficiency (km^2^/day) of unattended nests is defined by two components to include complete and partial nest predation:

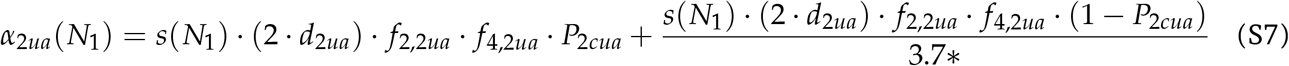

and handling time (day/per nest) of unattended nests is defined as:

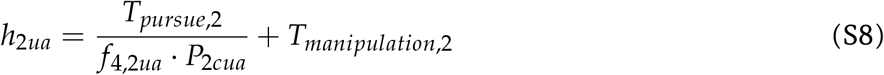

*This value refers to the average clutch size of the greater snow goose (Gauthier et al., 2013).

#### 7.2.2 Acquisition rate of attended goose nests by arctic foxes (prey 2)

The number of attended nests acquired is expressed as:

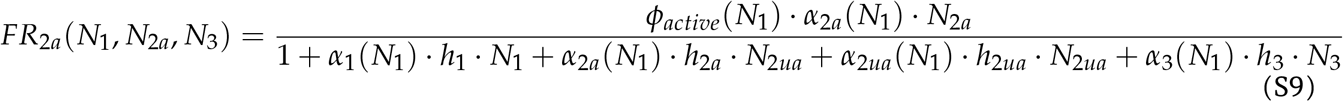

where density of attended goose nest is expressed as:

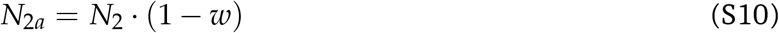

capture efficiency (km^2^/day) of attended nests is defined as:

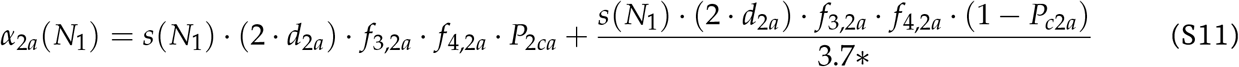

and handling time (day/per nest) of attended nests is defined as:

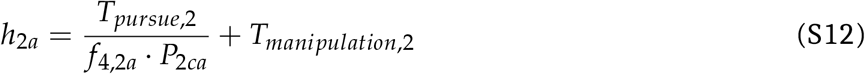

*This value refers to the average clutch size of the greater snow goose (Gauthier et al., 2013).

### 7.3 Acquisition rate of sandpiper nests by arctic foxes (prey 3)

The number of sandpiper nests acquired per fox per day (the acquisition rate) is expressed as:

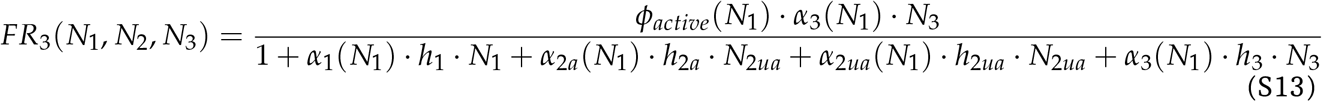

Where capture efficiency (km^2^/day) is defined as:

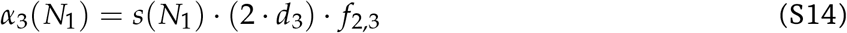

and handling time (day/per nest) is defined as:

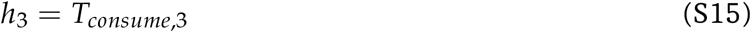

## Appendix S3 Details of the sandpiper population model

We used a projection matrix model developed by Weiser et al. (2020) for the semipalmated sandpiper but with a modification. As it is a post-breeding model, we added either adult or juvenile survival to the fecundity terms of the model. Using the the original model of Weiser et al. (2020) leads to similar results and the same conclusions.

The post-breeding projection matrix model is male-based and includes three age classes composed of juveniles, yearlings and 2+ years old (Fig. S1). Variations in the size (Z) of the age structured population between times t and t + 1 can be computed from:

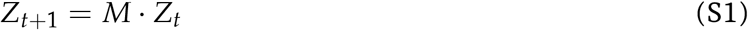

where M is a population projection matrix and Z is a vector describing the age-structured population. The projection matrix is:

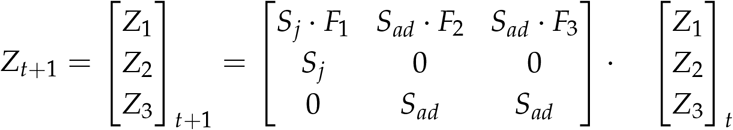

**Figure S1:**
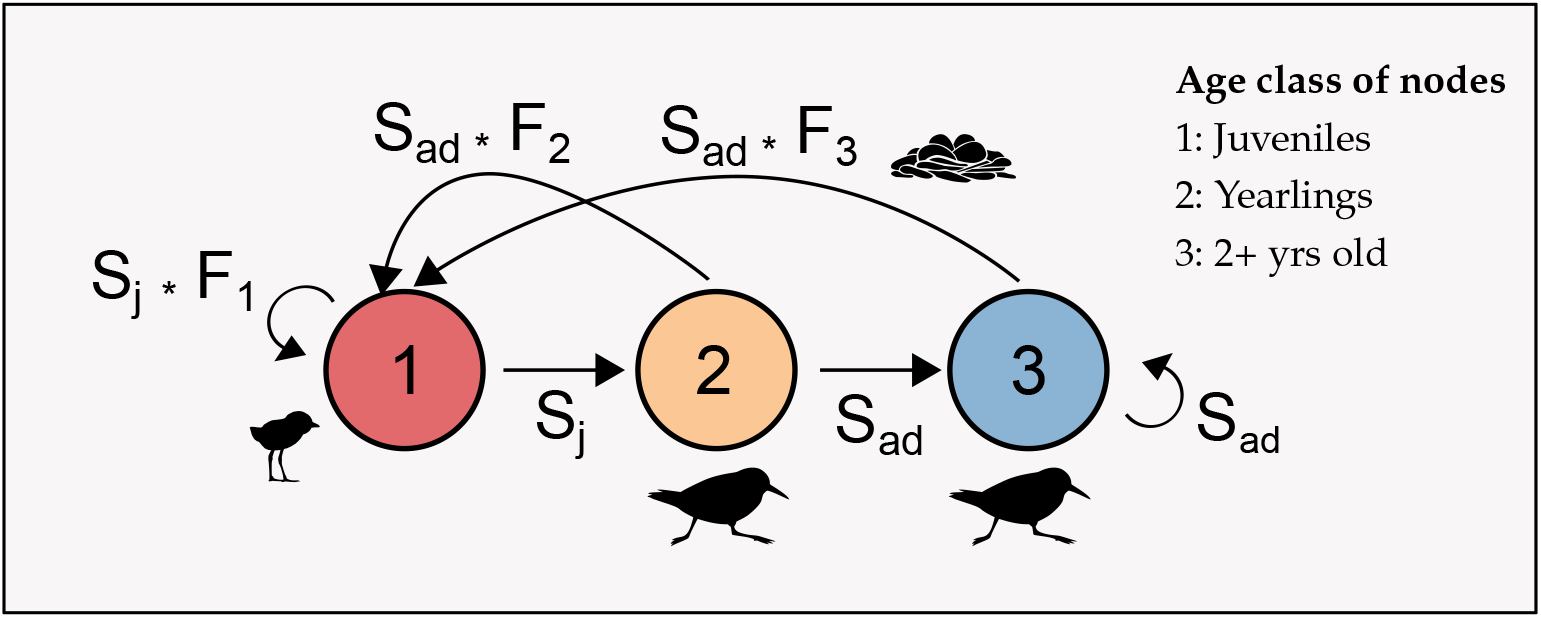
Diagram for the age-structured population model of semipalmated sandpiper.

Transitions among age classes are described by annual juvenile survival (*S_j_*) and adult survival (*S_ad_*). Age-specific probabilities of returning to the breeding area resulted in age-specific fecundity values (*F*_1_, *F*_2_, and *F*_3_), but survival values (*S_j_* and *S_ad_*) did not vary among classes after the first year because insufficient data were available to develop age-specific estimates. The fecundity value (F) associated with each age class is defined by the sum of *F_ini_* and *F_renest_* and the probability of returning to breeding area (e.g, *P*1_*return*_ for yearlings), which is essentially an age-specific recruitment probability, as follows:

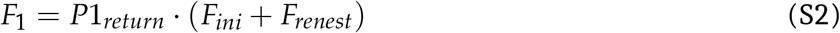

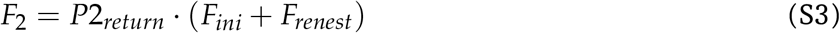

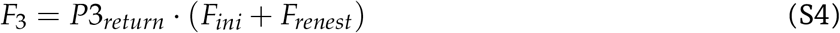

The mean number of male fledglings produced per breeding adult male for initial nests (*F_ini_*) and renesting attempt (*F_renest_*) is calculated as follows:

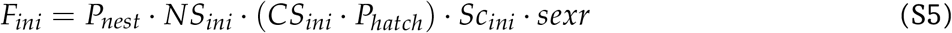

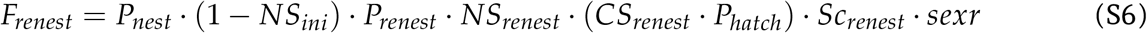

where fecundity of initial nests is defined by the probability of nesting in a given year (*P_nest_*), the average nesting success (*NS_ini_*), the number of eggs expected to hatch (the average clutch size, *CS_ini_*, multiplied by the hatching probability, *P_hatch_*), the survival rate of chicks to fledging (*SC_ini_*), and the sex ratio (the proportion of eggs that were expected to be male, *sexr*). The fecundity of renesting birds was conditional on the failure of the initial nest (1 – *NS_ini_*). See Table S1 for a complete description of the model parameters.

**Table S1:** Parameters used in the population matrix model of semipalmated sandpiper. Details regarding the estimation of parameter values can be found in Weiser et al. (2020).

| Parameter name | Symbol | Value (sd) |
| --- | --- | --- |
| Probability of returning to breeding area (yearlings) | $P1_{return}$ | 0.67 (0.10) |
| Probability of returning to breeding area (age 2) | $P2_{return}$ | 0.925 (0.10) |
| Probability of returning to breeding area (age 3+) | $P3_{return}$ | 1 |
| Probability of nesting in a given year | $P_{nest}$ | 0.8-1* |
| Mean clutch size (initial nests) | $CS_{ini}$ | 3.89 |
| Mean clutch size (renests) | $CS_{renest}$ | 3.76 |
| Sex ratio of eggs | $sexr$ | 0.5 |
| Average nesting success probability (initial nests) | $NS_{ini}$ | Values from the predation model |
| Average nesting success probability (renests) | $NS_{renest}$ | Values from the predation model |
| Proportion of eggs that hatch | $P_{hatch}$ | 0.94 (0.01) |
| Probability of renesting in a given year | $P_{renest}$ | 0.73 (0.20) |
| Chick survival probability (initial nests) | $SC_{ini}$ | 0.71 (0.07) |
| Chick survival probability (renests) | $SC_{renest}$ | 0.23 (0.19) |
| Juvenile survival probability | $S_j$ | 0.44 (0.10) |
| Adult (1+ yr old) survival probability | $S_{ad}$ | 0.76 (0.09) |

\* Since empirical data of nesting probability are virtually absent (Weiser et al., 2020), we ran simulations using nesting probability values ranging from 0.8 to 1.

We calculated growth rate (*λ*) using the mean values of each vital rates (Table S1) and average nesting success value obtained from different predation models. Given the strong influence of annual adult survival on *λ* and since empirical data of nesting probability are virtually absent (Weiser et al., 2020), we conducted simulations for a range of value of adult survival (0.76 ± 5%) and nesting probability (from 0.8 to 1).

